# Fire ! The spread of the Caribbean fire coral *Millepora alcicornis* in the Eastern Atlantic

**DOI:** 10.1101/519041

**Authors:** Peter Wirtz, Carla Zilberberg

**Affiliations:** Centro de Ciências do Mar, Universidade do Algarve, Campus de Gambelas, PT 8005-139 Faro, Portugal.; Núcleo em Ecologia e Desenvolvimento Ambiental de Macaé, Universidade Federal do Rio de Janeiro, Macaé, Brazil.

**Keywords:** Long-distance dispersal, Millepora alcicornis, phylogeography, habitat change

## Abstract

The Western Atlantic fire coral *Millepora alcicornis* is recorded from Madeira Island in the Eastern Atlantic for the first time. A large colony of this species has apparently been present at a very exposed site at the northern shore of Madeira Island for more than 15 years. Genetic analyses suggest that the colonies of this tropical fire coral at a mid-Atlantic location (Ascension Island) and at each of three Eastern Atlantic locations (Cape Verde Islands, Canary Islands, Madeira Island) originated from independent long-distance dispersal events from the Caribbean area.

## INTRODUCTION

Long-distance dispersal events occur rarely but, lying at the heart of island biogeography theory, they play a fundamental role in shaping species large-scale biogeography (Smith et al. 2018). The arrival of a species into a new area can have profound impacts on local ecological communities, leading altered environmental conditions and novel interactions, in particular if that species is a habitat forming one and competing with local endemics.

Fire corals (*Millepora* species) are well known for being important reef builders because of their large calcareous skeletons and for inflicting painful stings to humans (Lewis 2006). The genus is limited to 50 m depth in tropical seas, with a clear distinction between Atlantic and Pacific species (Razak & Hoeksema 2003). There are seven species in the Atlantic Ocean and until recently these were only reported from the tropical western Atlantic and from the Cape Verde Islands in the Eastern Atlantic (Laborel 1974, de Weerdt 1984). The species found at the Cape Verde Islands, *Millepora alcicornis* Linnaeus, 1758, is common in the western Atlantic and has recently also been documented from Ascension Island in the middle of the Atlantic Ocean (Hoeksema et al. 2017) and from the Canary Islands in the Eastern Atlantic, where it apparently arrived only a short time ago (Clemente et al. 2011, López et al. 2017).

In summer 2016, a fisherman from Madeira Island showed a large dried branch of a fire coral (*Millepora* sp.) to the first author and claimed that it came from the north coast of this island. The previous finding of fire coral at the Canary Islands (28°N) was already quite unexpected; the presence of *Millepora* in the even colder waters of Madeira Island (32°N) would be an even greater surprise.

Here we report that there is indeed a large colony of fire coral at Madeira’s north coast, the type of habitat it occupies, the identity of the species, and the probable origin of the colony.

## MATERIAL AND METHODS

The *Millepora* colony was photographed under water and several branches were collected in August 2016. Some of these branches are now deposited in the Natural History Museum of Funchal, Madeira, with the registration number MMF 46310. Fragments were preserved in 96% ethanol and sent to the second author for molecular analyses.

Total DNA extraction followed the phenol-chlorophorm method performed by Fukami et al. (2004), placing the sample in a CHAOS solution one week prior to extraction. DNA quality and concentration were assessed on a 0.8% agarose gel stained with GelRed (Biotium) and visualized under UV light, using the pattern Lambda DNA (125 ng/μL).

For species identification, the 16S rRNA gene of mitochondrial DNA (16S) was obtained from the Madeira *Millepora* colony and compared with previously reported sequences from other Atlantic *Millepora* specimens and species from the NCBI database (https://www.ncbi.nlm.nih.gov). A 537bp fragment of the large ribosomal subunit of the mitochondrial DNA (16S) was amplified using the following pair of primers: SHA 5'-ACGGAATGAACTCAAATCATGT-3; SHB 5'-TCGACTGTTTACCAAAAACATA-3’ (Cunningham & Buss 1993).

The polymerase chain reactions (PCR) consisted of PCR buffer 1X, dNTP (2 mM), bovine serum albumin (1 mg/ml), MgCl2 (1.5 mM), Taq polymerase (1U), primers (0.5 uM), ~1ng of template DNA. Thermal cycling conditions started with a denaturing step at 95°C for 3 min, followed by 10 cycles of 94°C for 1min, 40°C for 1min and 72°C for 1min, 40 cycles at 94°C for 1min, 52°C for 1min and 72°C for 1min and a final extension step at 72°C for 5 min. The amplified product was purified with ExoSAP-IT PCR Product Cleanup (Thermo Fisher Scientific) following manufacturer’s instructions and Sanger sequencing was performed in both directions at GATC Biotech (Germany).

Electropherograms were edited and a consensus sequence was created with Geneious R7 (http://www.geneious.com, Kearse et al., 2012). Alignment was performed using the ClustalW package in Geneious R7. Maximum likelihood (ML) phylogenetic reconstruction analyses and substitution models’ calculations were performed with PhyML 3.0 (Guidon et al. 2010). Substitution model selection was calculated using Smart Model Selection (Lefort et al. 2017) with the Akaike Information Criterion and the ML reconstruction. The substitution model used for the phylogenetic tree reconstruction was the HKY85 +G+I. Maximum likelihood analysis started with a neighbour joining tree followed by a Nearest Neighbour Interchange searching criterion and 1000 bootstraps for branch support.

A median-joining haplotype network was constructed using the software Networ v4.6.1.1 (Fluxus Technology Ltd.). This haplotype network included all *M. alcicornis* sequences used in de Souza et al. (2017), in addition to the sequence generated from the *Millepora* sample from Madeira Island in this study: Forty-four *Millepora alcicornis* colonies from the Caribbean Province, 109 colonies from the Brazilian Province, nine colonies from the Cape Verde Islands, a single colony from the Canary Islands, and two colonies from Ascension Island (Hoeksema et al. 2017) were compared with the colony from Madeira Island (Table 1).

**Table 1.**
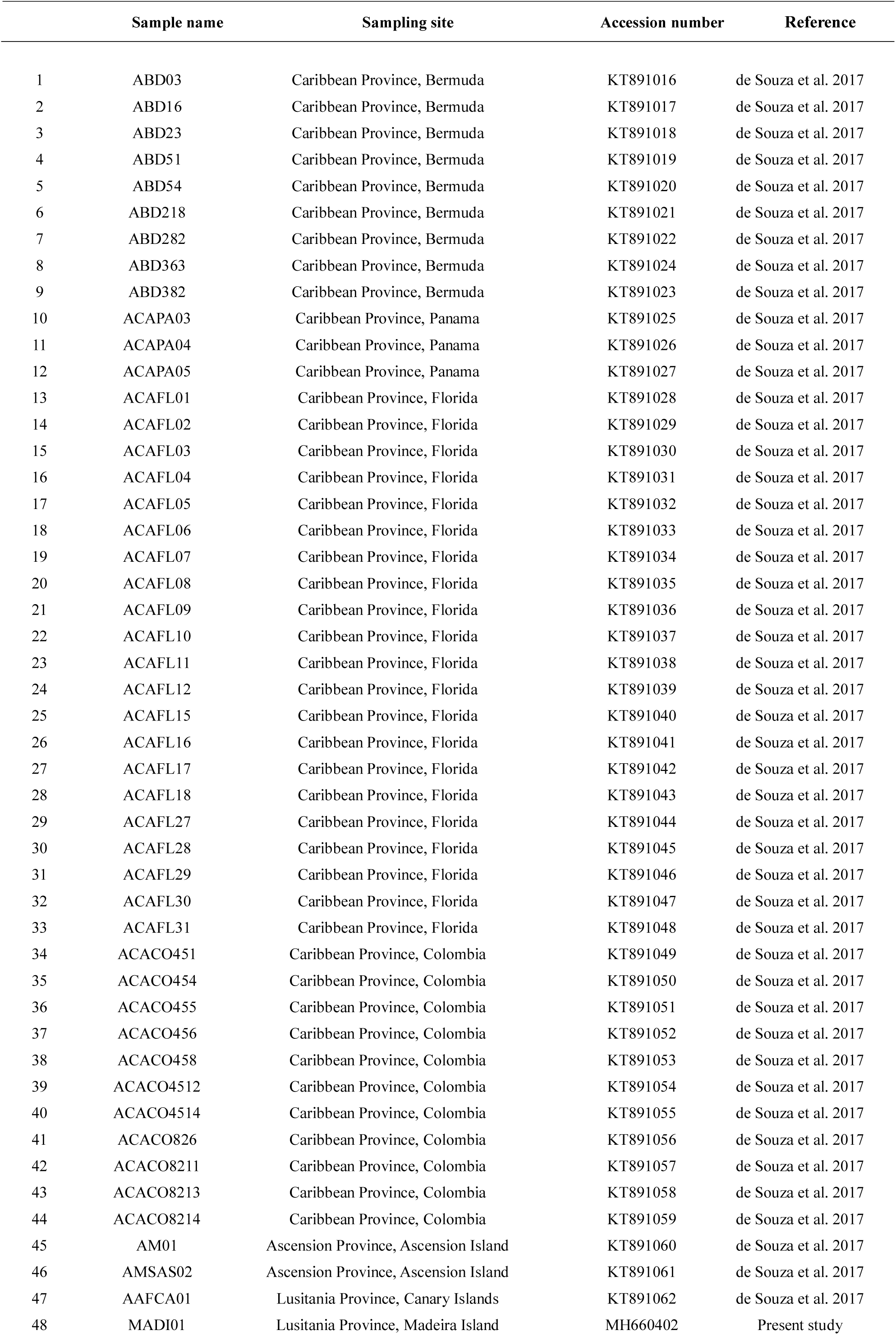
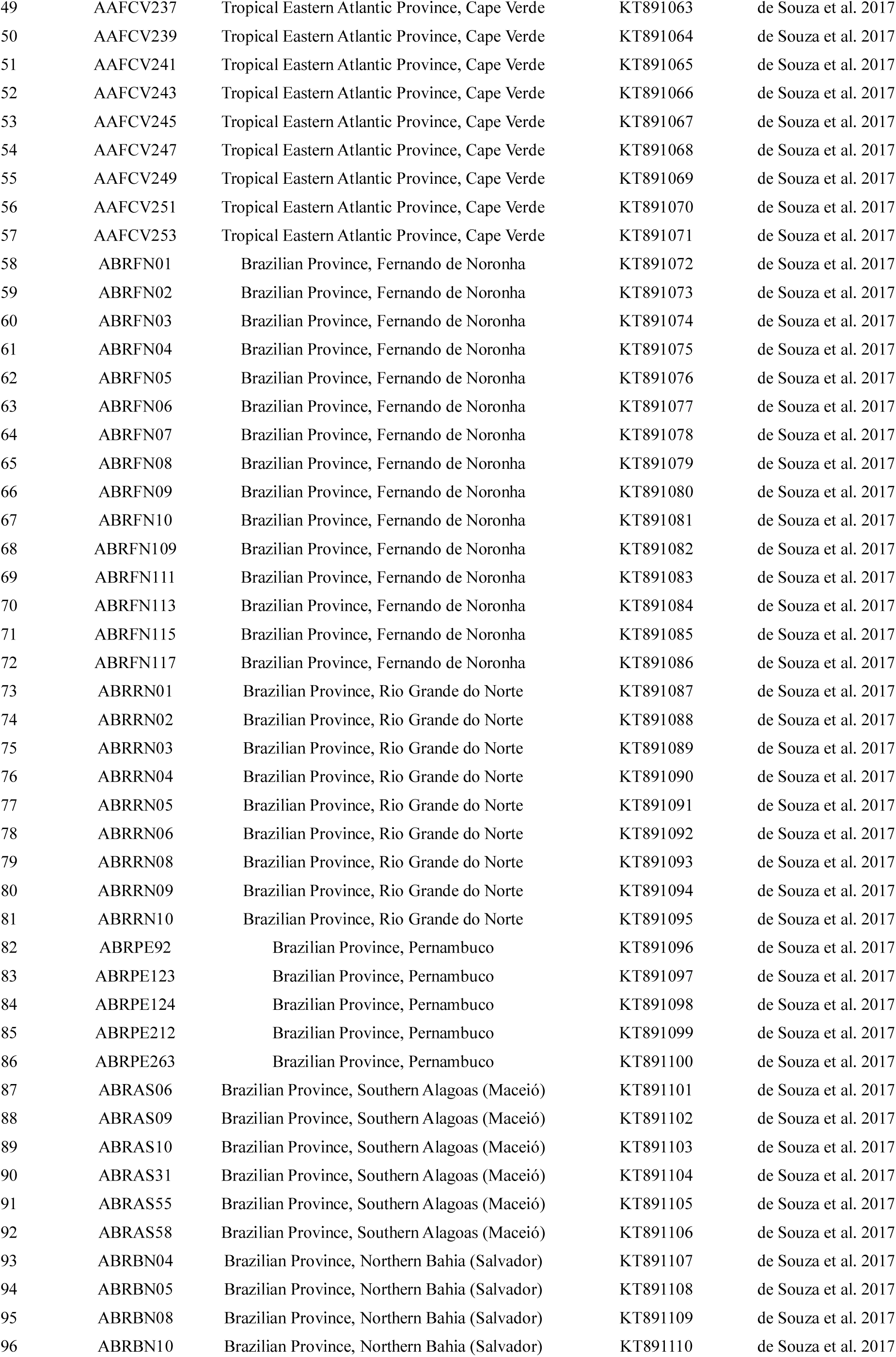
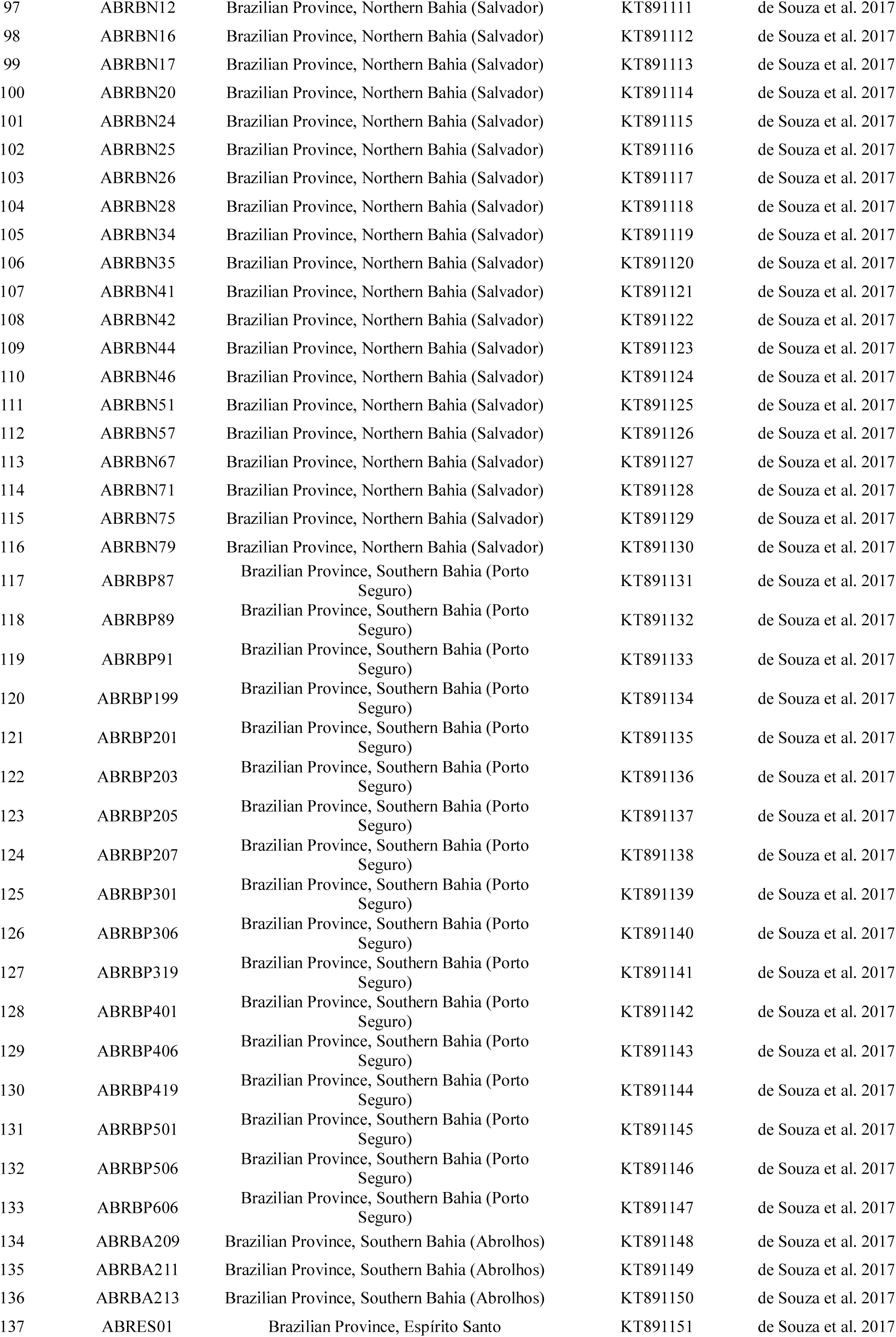
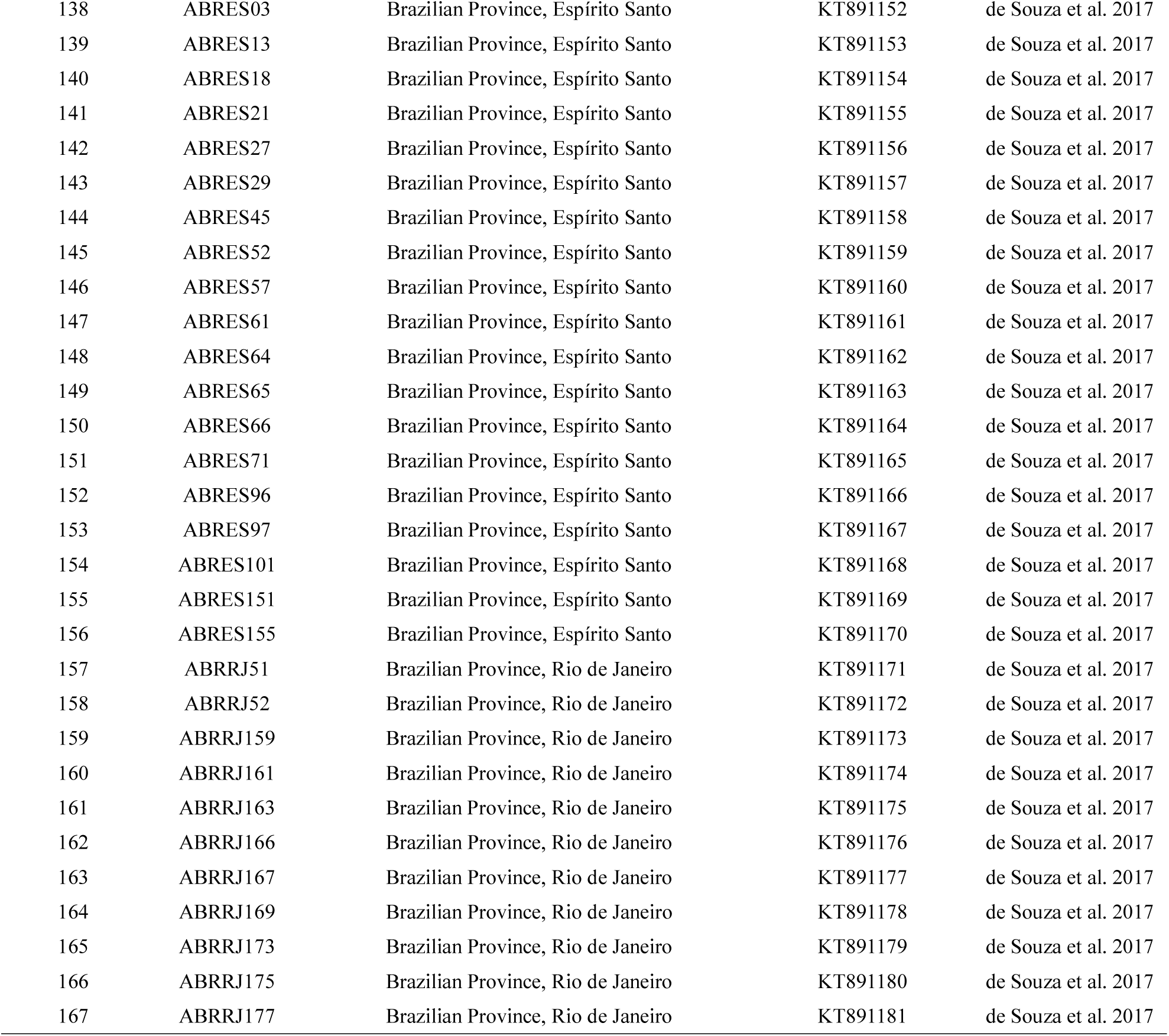
List of the16S rRNA sequences of Millepora alcicornis used in this study with their respective sampling sites and GenBank accession numbers.

## RESULTS

### Colony site and colony morphology

The *Millepora* colony of Madeira Island was found in shallow water (3m depth at low tide), in a small bay at a very exposed site on the north coast of Madeira Island (approximately 32°45′N, 16 o43′E). Water temperature at this site varies from 16 to 23 degrees C annually (personal observations by the first author).

The main colony had a roughly rectangular shape, approximately 4 m long and 3 m wide (figure 1). There were numerous small colonies (hand-sized or smaller) scattered around the large main colony. The fisherman, who guided the first author to this site, reported that there were fewer small colonies previously but that the main colony was already this large size when he first encountered it more than 15 years ago.

**Figure 1:**
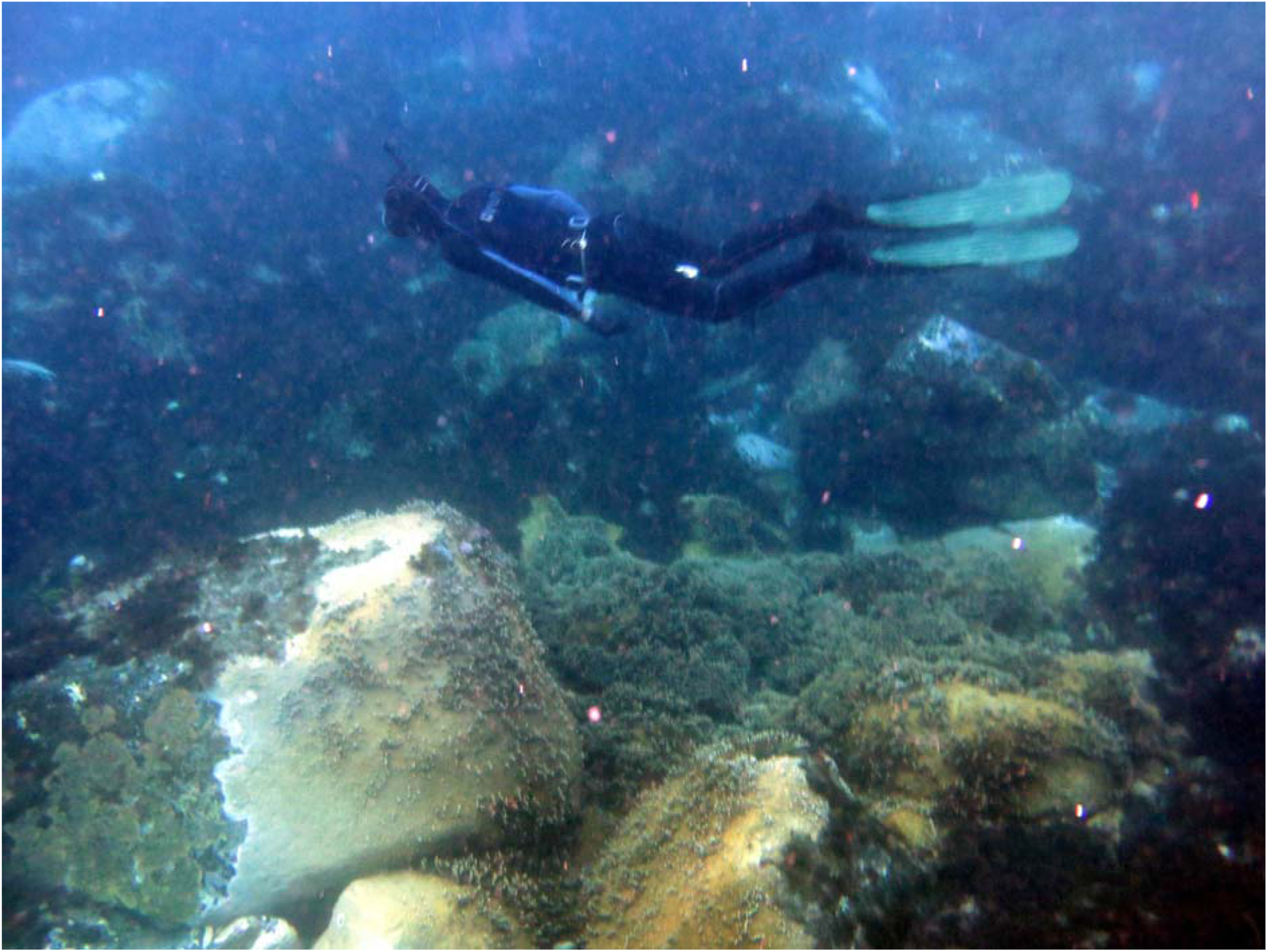
*Millepora* colony at the north coast of Madeira Island. The fins of the diver are 80 cm long.

The central part of the colony was characterized by erect branches up to 18 cm high, flattened laterally at the tips (Figures 2-3); at the edges and at its base, the colony was encrusting. The strong branches were very solid and difficult to break off, being able to resist the heavy wave action typical for the north coast of Madeira Island.

**Fig 2:**
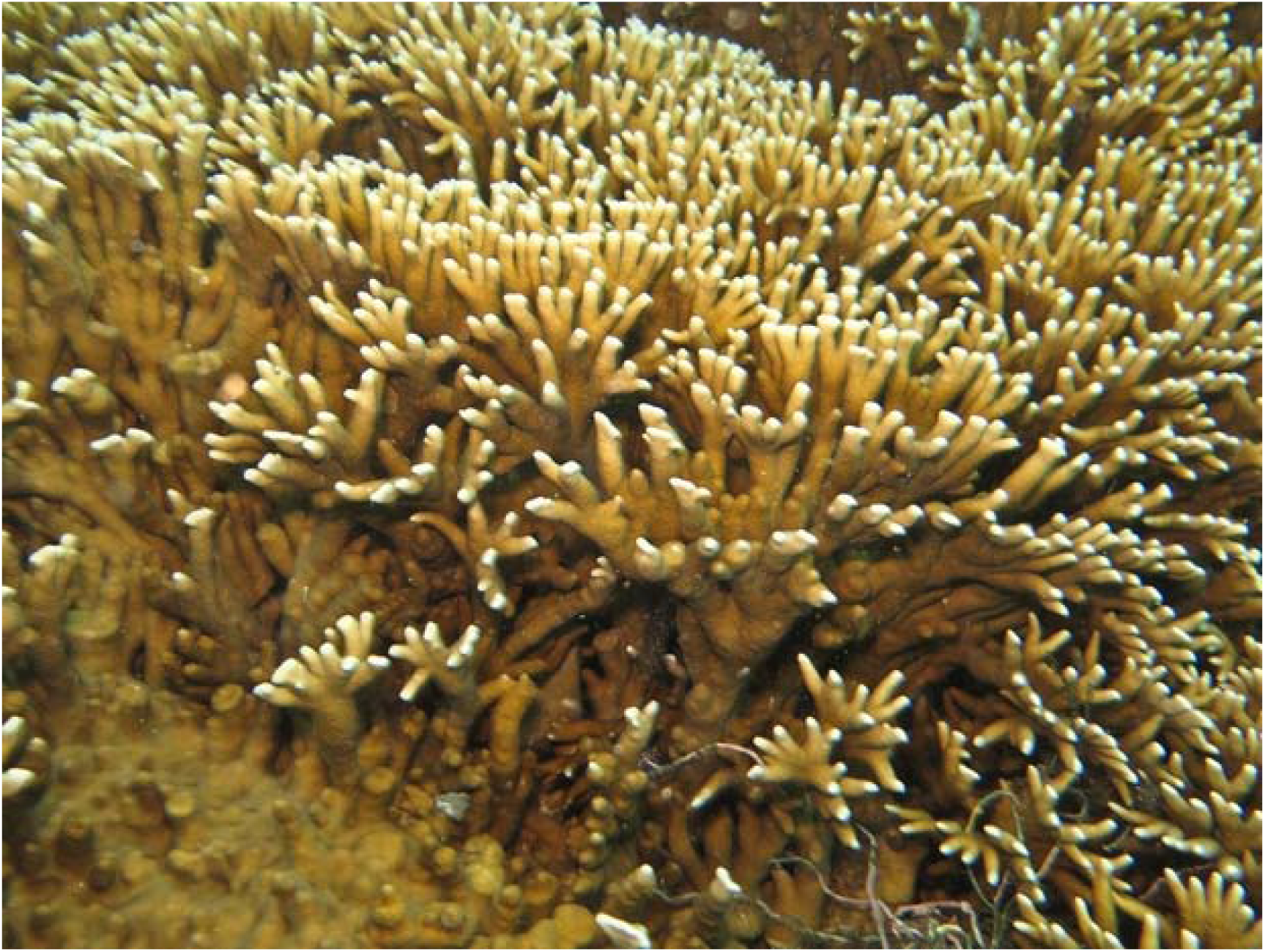
Detail of the main colony, showing the branching pattern and the erect branches typical of *M. alcicornis*, with laterally flattened tips.

**Figure 3:**
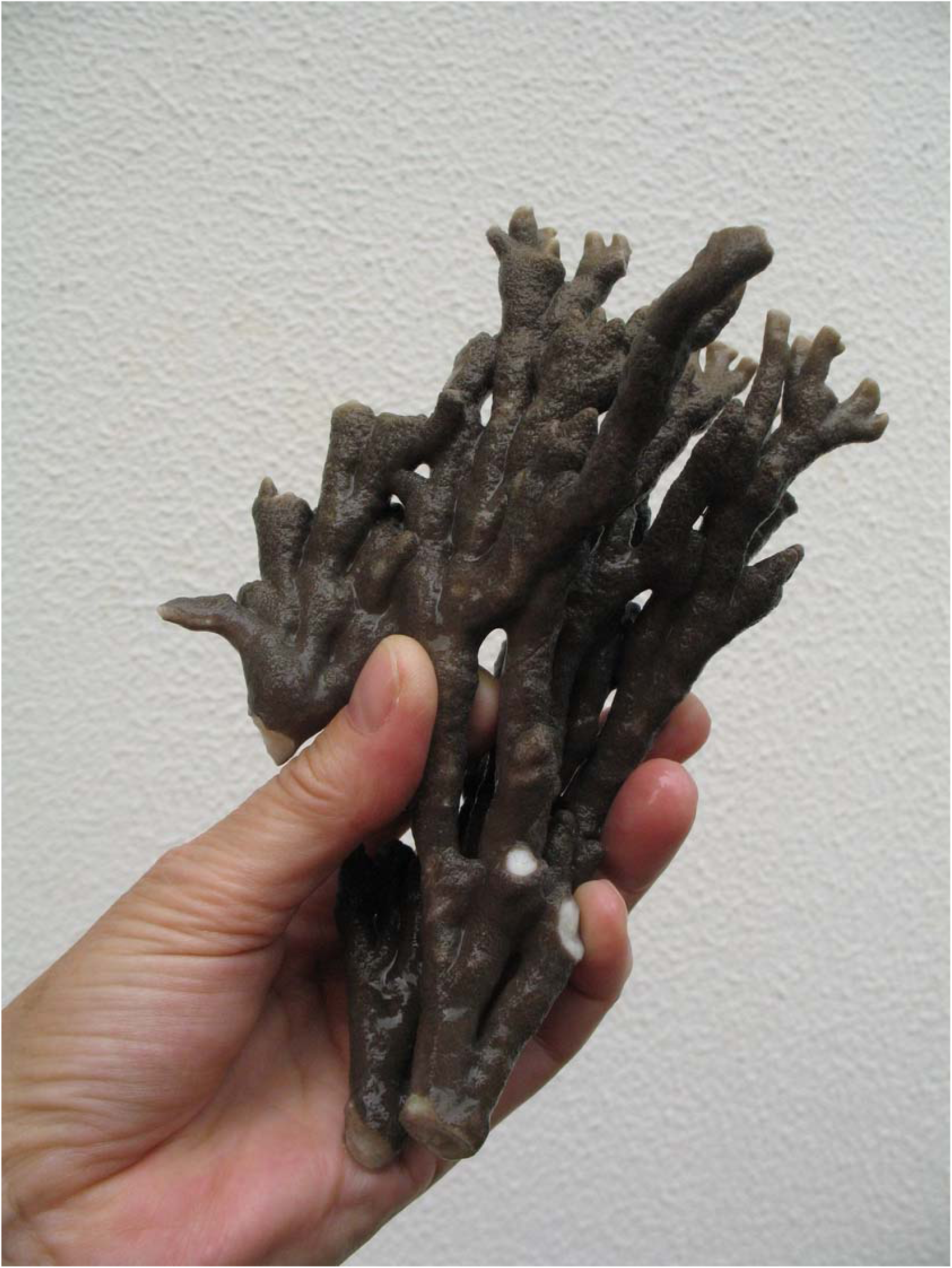
16 cm high branch of the main colony.

The growth form of the Madeiran *Millepora* colony was typical of - but not restricted to - the species *Millepora alcicornis* Linnaeus, 1748 (de Weerdt 1984, Hoeksema et al. 2017).

### Genetic analyses

#### a) Species identity

The phylogenetic analysis of the 16S sequences recovered the Madeira colony as *M. alcicornis*, with 89 % of bootstrap support, confirming the preliminary species identification made according to colony morphology (Figure 4).

**Figure 4.**
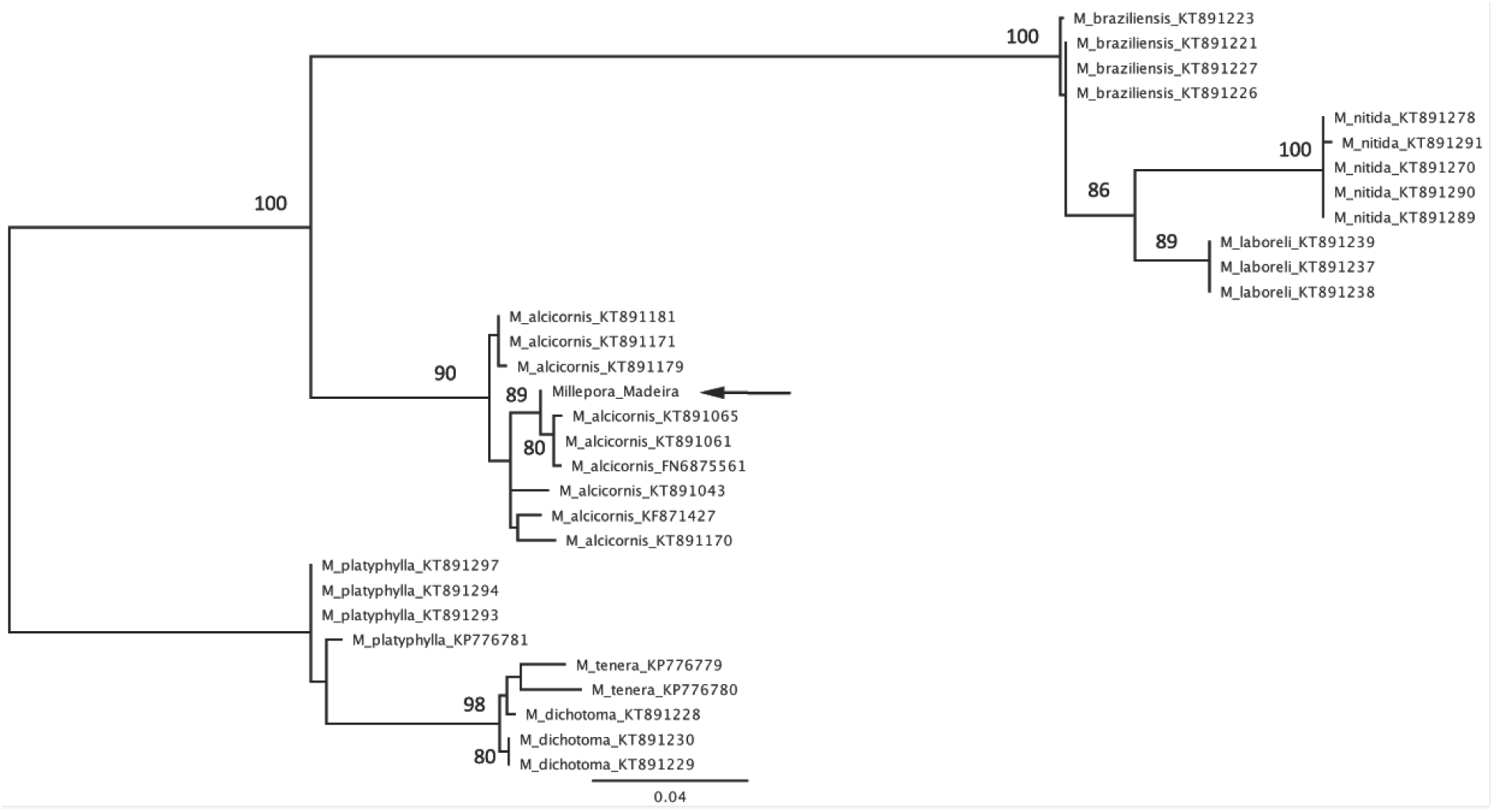
Maximum likelihood phylogenetic reconstruction of the partial 16S rRNA region of *Millepora* species. An arrow shows the sample from Madeira Island, grouping with *M. alcicornis*. Numbers at nodes represent the bootstrap support values (> 60%). All sequences downloaded from GenBank are presented with their access number. Genbank number of Madeira sample is MH660402.

#### b) Relationship of the colony

Figure 5 shows the haplotype network of the 16S fragment analyzed. Forty-three different haplotypes were found. The haplotype of the Madeiran colony corresponds to a Caribbean haplotype (with identical haplotype recorded at Bermuda and Florida) but not to a haplotype recorded from any of the other areas. This haplotype is only 2bp different from the Ascension Island haplotype.

**Figure 5.**
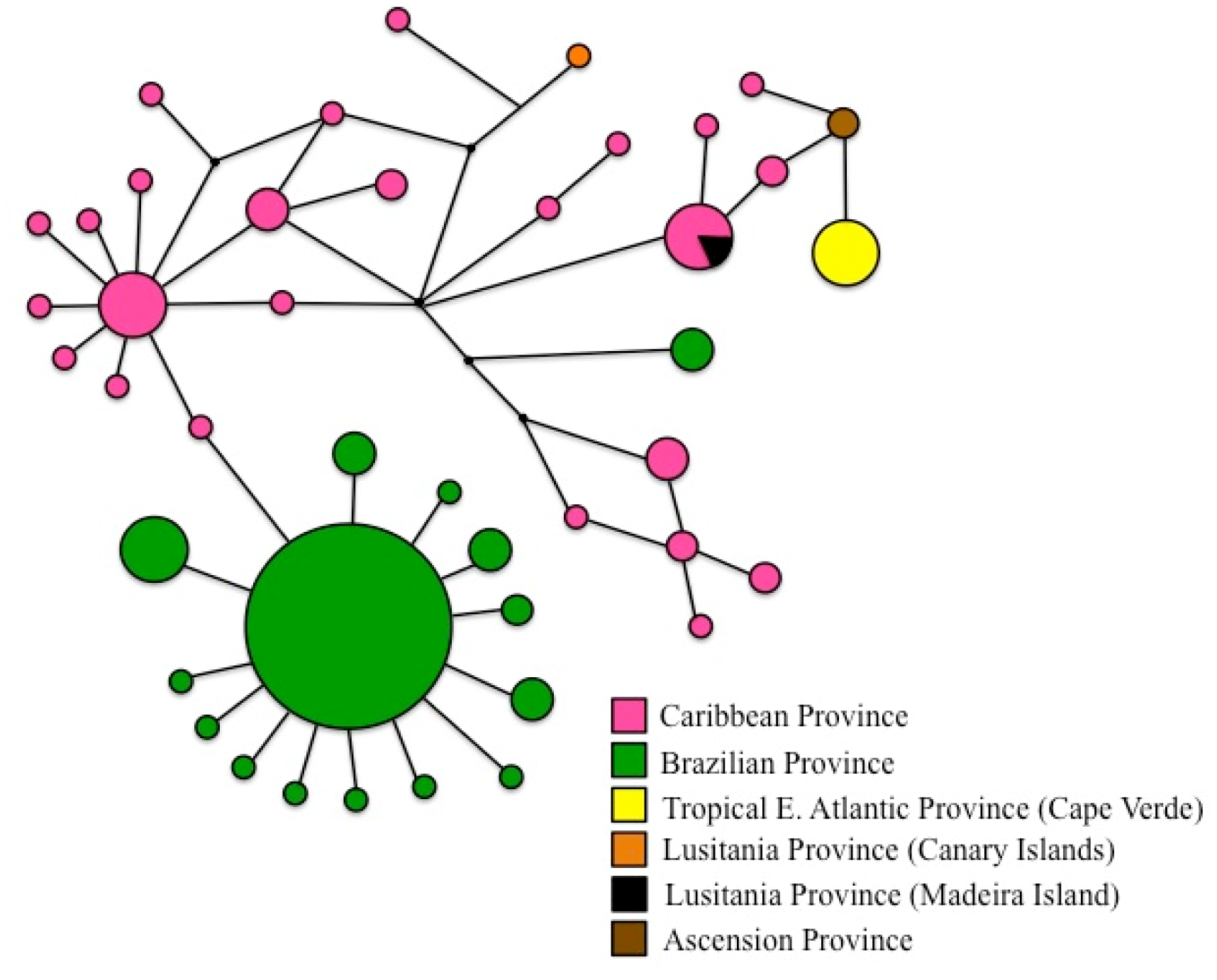
Haplotype network of *Millepora alcicornis* using sequences from de Souza et al. (2017; Table 1), including the *M. alcicornis* sample from Madeira Island. Circle sizes correspond to the number of haplotypes and colours illustrate major regions of occurrence. The four black dots represent ancestral or not sampled haplotypes and line length corresponds to distance between haplotypes, the smallest distance indicating one mutational event.

## DISCUSSION

In the Western Atlantic, *Millepora* species are most abundant in shallow, turbulent environments (Lewis 1989), i.e. in places similar (in these aspects) to the site where *Millepora alcicornis* has now been found at Madeira Island. To find the tropical fire coral *Millepora alcicornis* in the comparatively cold waters of Madeira Island (32°N) comes as a surprise and extends the known range of the species in the Eastern Atlantic about 550 km to the north.

In the Western Atlantic, the northern limit of *M. alcicornis* is at the Bermuda Islands, i.e. also at 32°N, but in a location with considerably warmer water temperatures than those at Madeira Island, winter minimum temperature there being 20°C (Bertelsen & Ussing 1936); this is four degrees warmer than at the north coast of Madeira Island. Thus, the Madeiran colony of *M. alcicornis* is living in the coldest environment ever reported for this species and this may well be the limit of the temperature tolerance of the species.

It is tempting to ascribe the recent discovery of a tropical species outside its known range to the effects of global warming (cf. Hansen et al. 2010). While this may be true for the new and rapidly growing colony at the Canary Islands (Clemente et al. 2011, López et al. 2015), this is a less likely explanation for the presence of fire coral at Madeira Island. The main colony already was large (about 4×3m area) when detected more than 15 years ago. Clemente et al. (2011) and González (2017) report on the growth rate of the colony at the Canary Islands. In eight years, this colony increased from 1.100 cm^2^ to 34.704 cm^2^. Assuming a linear growth rate, this would mean that the Madeiran colony (about 120.000 cm^2^) would have an age of about 28 years; however, the growth rate of older colonies is likely to be lower than that of younger colonies and growth in the colder waters of Madeira is likely to be slower than at the Canary Islands. Therefore, the *Millepora alcicornis* colony at the north coast of Madeira is probably considerably more than 30 years old. The appearance of *Millepora* at Madeira Island is thus unlikely to have been facilitated by Global Warming.

Successful settlement is limited not only by favourable conditions on arrival; it is limited first of all by the probability of arrival. Genetic analysis suggested that the Madeiran *M. alcicornis* colony originated in a long distance dispersal event directly from the Caribbean area. Additionally, genetic analysis already suggested that the colonies of this tropical fire coral species at the mid-Atlantic location (Ascension Island) and at the Cape Verde Island, and at the Canary Island have also originated from settlement events from the Caribbean area (López et al. 2015, Hoeksema et al. 2017, de Souza et al. 2017). As the Madeira colony is only 2bp different from the Ascension Island colony, there remains a small possibility that it may have originated from there.

To date, no other marine species at Madeira Island has been shown to be directly related to the Caribbean area. The possible ways of arrival of *M. alcicornis* at the north coast of Madeira Island are natural larval dispersal, rafting of an adult colony, or transport by humans (in ballast water or attached to a ship hull). Human transport appears unlikely, as there are only small, local fishing harbours on the north coast of Madeira, and none of them near the colony: no ship from the Caribbean is likely to have passed there. Natural larval dispersal depends on currents. The North Equatorial Countercurrent directly connects the Caribbean and the Cape Verde archipelago. There are, however, no currents directly linking the Caribbean and Madeira Island. Madeira is bathed by a branch of the Gulf Stream, which first passes the Azores far in the north and then turns southward as the Canary Current. This would be a very long journey indeed for a *Millepora* larva; given the short life span of the medusa stage of *Millepora* (Lewis 2006), larval transport from the Caribbean to Madeira therefore also appears unlikely. *M. alcicornis* is capable of growing on artificial substrates (de Souza et al. 2017) and *Millepora* species have been reported to raft on ship hulls (Bertelsen and Ussing 1936) and pumice (Jokiel 1989). Transatlantic rafting from the Western Atlantic Ocean - as demonstrated already for various mollusc species (Holmes et al 2015) and the coral *Astrangia poculata* (Hoeksema et al. 2012, 2015) - appears to be the most likely explanation for the origin of the Madeira *Millepora* colony.

Long distance dispersal – long considered to be extremely rare (de Queiroz 2005) – is more common than previously thought and may even become more common under climate change conditions (Batista et al. 2018).

## ACKNOWLEDGEMENTS

Many thanks to the fisherman, who showed the colony to the first author and described its history; he wishes to remain anonymous. For helpful comments on an early draft of the manuscript, we are grateful to C. Lopez. This study received Portuguese national funds through FCT - Foundation for Science and Technology - through project UID/Multi/04326/2013.

